# CellSpliceNet: Interpretable Multimodal Modeling of Alternative Splicing Across Neurons in *C. elegans*

**DOI:** 10.1101/2025.06.22.660966

**Authors:** Arman Afrasiyabi, Jake Kovalic, Chen Liu, Egbert Castro, Alexis Weinreb, Erdem Varol, David M. Miller, Marc Hammarlund, Smita Krishnaswamy

## Abstract

Alternative splicing profoundly diversifies the transcriptome and proteome, but decoding its regulatory mechanisms remains a challenge. We introduce `CellSpliceNet`, an interpretable transformer-based multimodal deep learning framework designed to predict splicing outcomes across the neurons of *C. elegans*. By integrating four complementary data modalities, namely long-range genomic sequence, local regions of interest (ROIs) in the RNA sequence, secondary structure, and gene expression, `CellSpliceNet` captures the complex interplay of factors that influence splicing decisions within the cellular context. `CellSpliceNet` employs modality-specific transformer embeddings, incorporating structural representations guided by mutual information and scattering graph embeddings. To this end, a novel and carefully designed multimodal multi-head attention mechanism preserves the integrity of each modality while facilitating selective cross-modal interactions, notably allowing gene expression data to inform sequence and structural predictions. Attention-based pooling within each modality highlights biologically critical elements, such as canonical intron–exon splice boundaries and accessible single-stranded RNA loop structures within the exon. Quantitative comparisons with current state-of-the-art methods demonstrated `CellSpliceNet`‘s superior predictive accuracy (Spearman *ρ* = 0.88) and high accuracy across diverse neuron subtypes. Furthermore, `CellSpliceNet` elucidates a hierarchical, neuron-specific splicing code by preferentially weighting upstream enhancer motifs (e.g., GGAAGAAC) and identifying neuron-class-specific splicing-factor signatures, including *smu-1, unc-75*, and *hrp-1*. Thus, `CellSpliceNet` not only advances the frontiers of alternative splicing predictive capabilities but also provides mechanistic insights into the multimodal regulation of alternative splicing.

## Main

RNA alternative splicing is a fundamental mechanism that promotes functional diversity in eukaryotic organisms by allowing a single gene to produce multiple mRNA and protein isoforms [1], thereby greatly expanding the functional repertoire encoded by the genome [2]. More than 95% of human genes undergo alternative splicing [2, 3], giving rise to distinct variants that help regulate cellular processes such as differentiation [4]. Disruptions to this finely tuned mechanism are known to be associated with a wide range of diseases, including cancer [5], neurodegenerative disorders [6], and developmental abnormalities [7], underscoring the biological importance of alternative splicing.

Despite its significance, computationally modeling RNA alternative splicing remains difficult. Recent advances in biological sequence modeling have been driven by transformers [8] and large language models (LLMs) [9], which learn co-occurrence patterns and conserved motifs from vast repositories of naturally occurring sequences [10, 11, 12]. These approaches have outperformed earlier methods in numerous tasks, including information embedding [13], structure modeling [12], and protein folding [14, 15, 16]. However, these methods often overlook key drivers of splicing: local regulatory motifs near-splice junctions, spatial organization imposed by RNA secondary structures, and cell-specific factors that directly influence exon inclusion or skipping. As a result, these models provide limited insight into the mechanisms underlying splicing variation.

Several challenges hinder robust modeling of alternative splicing. First, RNA has a small “vocabulary” composed of only four nucleotides, which renders standard tokenization strategies designed for larger vocabularies less suitable. Second, binding motifs might degenerate and exhibit a degree of ambiguity, leading to high false discovery rates in motif detection. Moreover, non-coding regions introduce further noise due to relaxed evolutionary constraints. Finally, capturing cell-specific regulatory influences is nontrivial, as splicing patterns can change in response to intricate networks of splice factors and other regulatory proteins.

To address these challenges, we propose `CellSpliceNet`, a deep multimodal transformer-based framework that integrates four key sources of information: the full-length RNA sequence, its generated secondary structure, the regions of interest (ROIs) around splice junctions, and single-cell gene expression profiles. By explicitly modeling sequences near splice junctions, where regulatory elements often reside, and incorporating structural information, `CellSpliceNet` captures both global and local features critical to splice-site selection. To capture cell-specific regulatory influences, we use single cell expression data by imputing missing values with MAGIC [17] and deriving splice factor coexpression graphs using pairwise mutual information. Rather than utilizing graph neural networks, these graphs are encoded via geometric scattering [18], a structure-preserving signal processing transformation shown to be useful in numerous applications.

`CellSpliceNet` introduces several innovations. First, it adopts a patch-wise tokenization scheme tailored to RNA’s small vocabulary. Second, it focuses on biologically relevant regions of interest (ROIs) to counter motif degeneracy, where many RNA-binding proteins tolerate specific nucleotide substitutions, so several related sequences all qualify as the same binding site. Third, it encodes cell-specific context using “cell state tokens” to capture cell-specific splicing differences. Finally, a learnable pooling mechanism fuses modality-specific features with learnable and interpretable weights, offering insight into the contribution of each input type.

`CellSpliceNet` combines state-of-the-art predictive power with fine-grained mechanistic insight. Ablation tests reveal that each modality contributes uniquely: removing any modality lowers the performance. Benchmark comparisons show that `CellSpliceNet` consistent outperforms existing methods across five leading predictors, with strong generalization across neuron types. The attention landscape reflects and extends known biology: it emphasizes intron–exon boundaries while flexibly adapting to neuro-specific cues, prioritizes single-stranded loops in exons over distal stems, and concentrates nearly 40% of ROI attention to the upstream exon where splicing enhancers reside. A trainable fusion layer then reconciles cross-modal evidence, amplifying ROI and expression signals, probing late-sequence patches, and selectively down-weighting canonical motifs to resolve ambiguity. These capabilities position `CellSpliceNet` as a powerful and interpretable tool for decoding the rules of RNA splicing across diverse cellular contexts.

## Results

### Architecture of the CellSpliceNet model

Alternative splicing increases the transcriptomic and proteomic diversity by enabling a single gene to produce multiple protein isoforms. On average, each human gene gives rise to five to seven distinct isoforms [19, 20, 21, 22], and the disruption of alternative splicing is central to numerous pathologies [5, 6]. Among the various splicing mechanisms, exon skipping (Fig. 1a) is the most prevalent, in which specific exons are retained or omitted during mRNA processing. The degree to which an exon is included is quantified by the *percentage spliced in (PSI)* metric [23], which reflects the fraction of transcripts that incorporate the exon. PSI values range from zero to one and vary between tissues, developmental stages, and environmental conditions, reflecting the precise regulation of splicing in different cellular contexts [7].

**Fig. 1:**
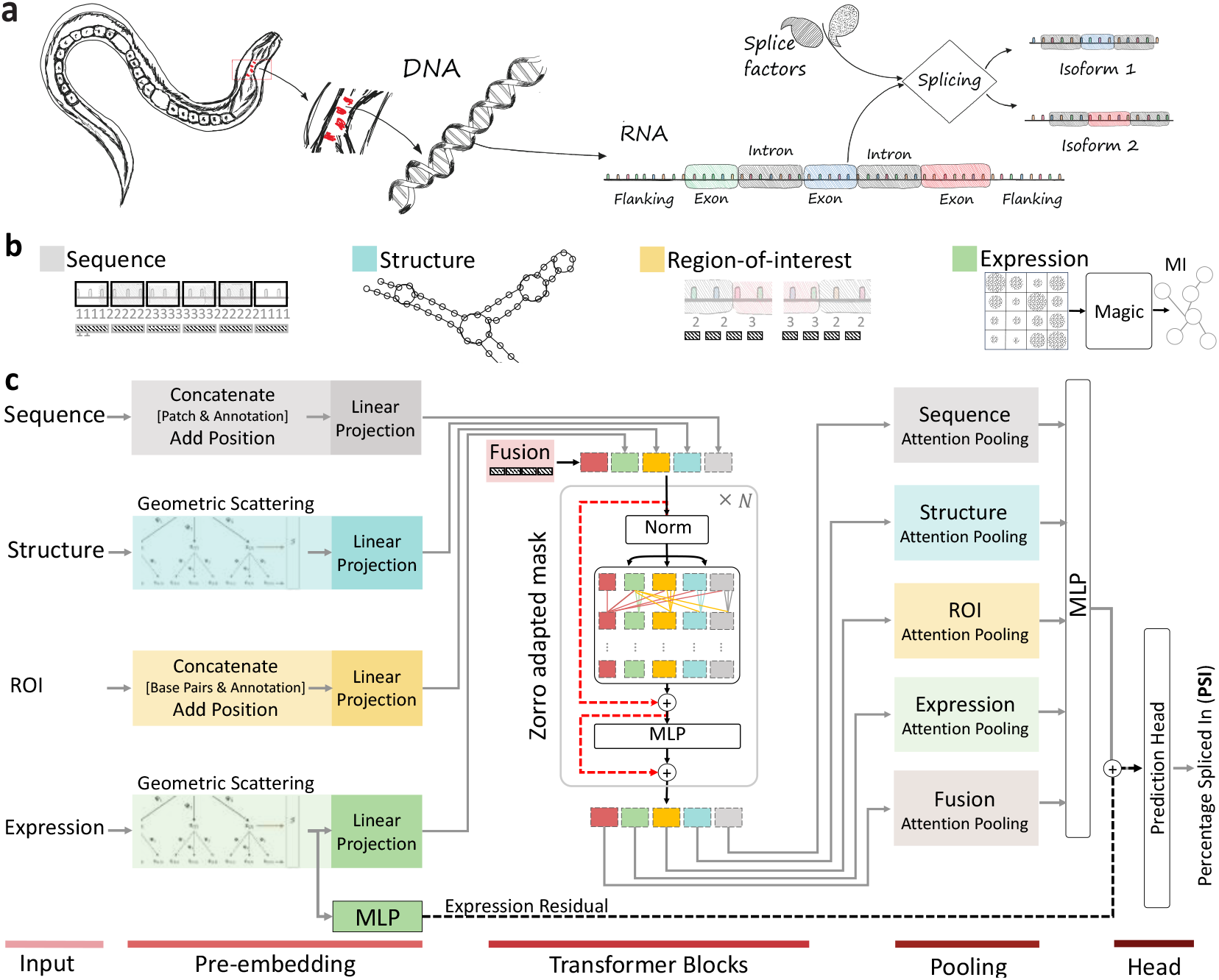
Multimodal CellSpliceNet for predicting alternative splicing. **a**. Schematic representation of alternative splicing in Caenorhabditis elegans. The illustration begins with the organism and narrows down to a single neuron, showcasing the detailed processes at the DNA and RNA levels, culminating in RNA processing. Splice factors are depicted as guiding the alternative splicing pathway, leading to the production of two distinct mRNA isoforms. **b**. Overview of the multimodal data integration approach, ranging from whole sequence analysis to secondary structure prediction, and region of interest (ROI) imputation using MAGIC [17] for graph construction based on mutual information (MI). **c**. Workflow of the `CellSpliceNet` model, processing inputs across local, global, and expression modalities (indicated by varying colors on the left) to predict the percentage spliced in (PSI), shown on the right side of the figure.

Recent studies have shown that the interaction between sequence motifs, secondary RNA structures, and the expression of splicing factors critically influences PSI outcomes [24, 25]. However, existing computational approaches typically model these factors in isolation, limiting their ability to capture the full regulatory complexity [26, 27, 28, 29].

To address this limitation, we developed `CellSpliceNet`, a multimodal transformer architecture designed to predict PSI from biologically grounded inputs. The model integrates four sources of information: (1) the full-length RNA sequence to capture long-range context, (2) the RNA secondary structure that may regions of interest (ROIs) obscure or reveal these motifs, (3) the regions of interest (ROIs) near each splice site where regulatory motifs are concentrated, and (4) the gene expression data conveying cell-specific splicing factor profiles. These inputs are encoded independently, then integrated through a fusion module to produce a unified representation for PSI prediction. The four modalities are illustrated in Fig. 1b, and how `CellSpliceNet` processes them to predict PSI is shown in Fig. 1c.

### Modeling the full-length RNA sequence

To capture long-range regulatory signals that affect splicing, we tokenize the full-length RNA sequence using a patch-based strategy. Using single nucleotides as tokens would confine a computational model to a shallow four-word vocabulary [13], providing little higher-order context; therefore, we instead group contiguous bases into overlapping k-mer “patches.” Each patch is projected into an embedding space through a linear layer, following the Vision Transformer paradigm [30]. Tokens are further annotated with their genomic context (exon, intron, or junction) to improve positional awareness. This representation allows the transformer encoder to model sequence-wide dependencies while retaining structural and spatial features of the gene body.

### Modeling the RNA secondary structure

RNA folding can influence exon inclusion by modulating the accessibility of splice sites and nearby regulatory motifs, a mechanism known as structural masking [31]. To model this, we extract the intronic regions flanking each target exon and predict their secondary structures using ViennaRNA [32]. These structures are converted into graphs, with nodes representing nucleotides and edges representing base pairs. We apply the geometric scattering transform [18] to these graphs, producing fixed-length embeddings that capture multi-scale spatial patterns. This structural modality enables the model to learn how RNA folding affects regulatory motif exposure and ultimately splicing decisions.

### Modeling the regions of interest

Regions near splice junctions are enriched for cis-regulatory elements such as enhancers and silencers that modulate exon inclusion. To isolate these signals, we define regions of interest (ROIs) that span 200 base pairs upstream and downstream of each splice junction. These local sequence windows are embedded independently from the full-length transcript to emphasize short-range sequence features critical to splicing. By separately encoding ROIs, the model focuses attention on the intron-to-exon and exon-to-intron boundaries where splicing signals are concentrated.

### Modeling the gene expression and splice factor dynamics

Splicing decisions vary across cell types due to differences in the expression and activity of splicing factors. To incorporate this regulatory context, we use single-cell RNA-seq data and impute missing values with MAGIC [17]. We focus on 243 curated splicing factors and construct expression matrices stratified by neuron type. To capture co-regulation, we compute pairwise mutual information between splicing factors and discretize these values to generate co-expression graphs. Nodes represent splicing factors, and edges reflect significant mutual dependencies. We encode these graphs using geometric scattering [18], using Dirac signals weighted by average expression. This yields a compact representation of the regulatory landscape influencing exon selection.

### Fusing the heterogeneous modalities

The four modalities are respectively processed by a dedicated transformer encoder. To allow selective information exchange while maintaining modality-specific learning, we apply an attention-masking strategy inspired by Zorro [33]. This mechanism restricts direct cross-modal attention except through designated fusion tokens. We permit the expression modality to influence all others, as splicing factors can affect both sequence and structural mechanisms. The encoded modality embeddings are concatenated and passed to a multilayer perceptron to predict PSI values. The model is trained end-to-end by minimizing the squared error loss, as described in Eqn (1).

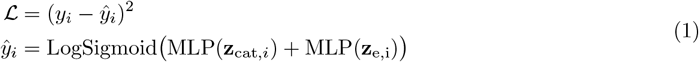

Here, **z**_cat,*i*_ represents the concatenated embeddings from all modalities and **z**_e,*i*_ is the residual expression component. This formulation ensures predictions remain within biologically meaningful bounds while penalizing large deviations from ground truth.

### Modality-specific interpretation mechanism

To enhance interpretability, `CellSpliceNet` employs attention-based pooling for each modality. After transformer encoding, token embeddings are aggregated using learnable attention weights following ReLSO [34]. Each pooling network computes a weighted sum of tokens within a modality, where weights are constrained to sum to one. This produces a single vector per modality that reflects its most informative features. In particular, the pooling distribution highlights biologically relevant patterns such as exon boundaries in sequence, structural motifs in folding, and high-variance regulators in expression. These vectors are concatenated and used for PSI prediction, providing an interpretable view of which modalities and features drive each prediction.

### CellSpliceNet improves splice inclusion prediction by integrating multimodal information

#### Four complementary modalities drive accurate splicing prediction

We first conducted a systematic ablation analysis to assess the individual and collective contributions of the four input modalities in `CellSpliceNet`, namely the full-length RNA sequence, region of interest (ROI), RNA secondary structure, and gene expression. As shown in Fig. 2a, the full multimodal model achieved the highest Spearman correlation of 0.88 in predicting PSI values. Removing any single modality led to a marked decline in performance (0.74 without the full sequence, 0.81 without ROI, 0.82 without structure, and 0.84 without expression), demonstrating the integrative value of these complementary signals. Columns 2-4 in Fig. 2a further illustrate the degradation in predictive accuracy via scatter plots comparing observed and predicted PSI values for models lacking the full sequence input (No Sequence), alongside the corresponding results from the complete model on both training and held-out test data. Together, these results highlight the necessity of multimodal integration for robust PSI prediction.

**Fig. 2:**
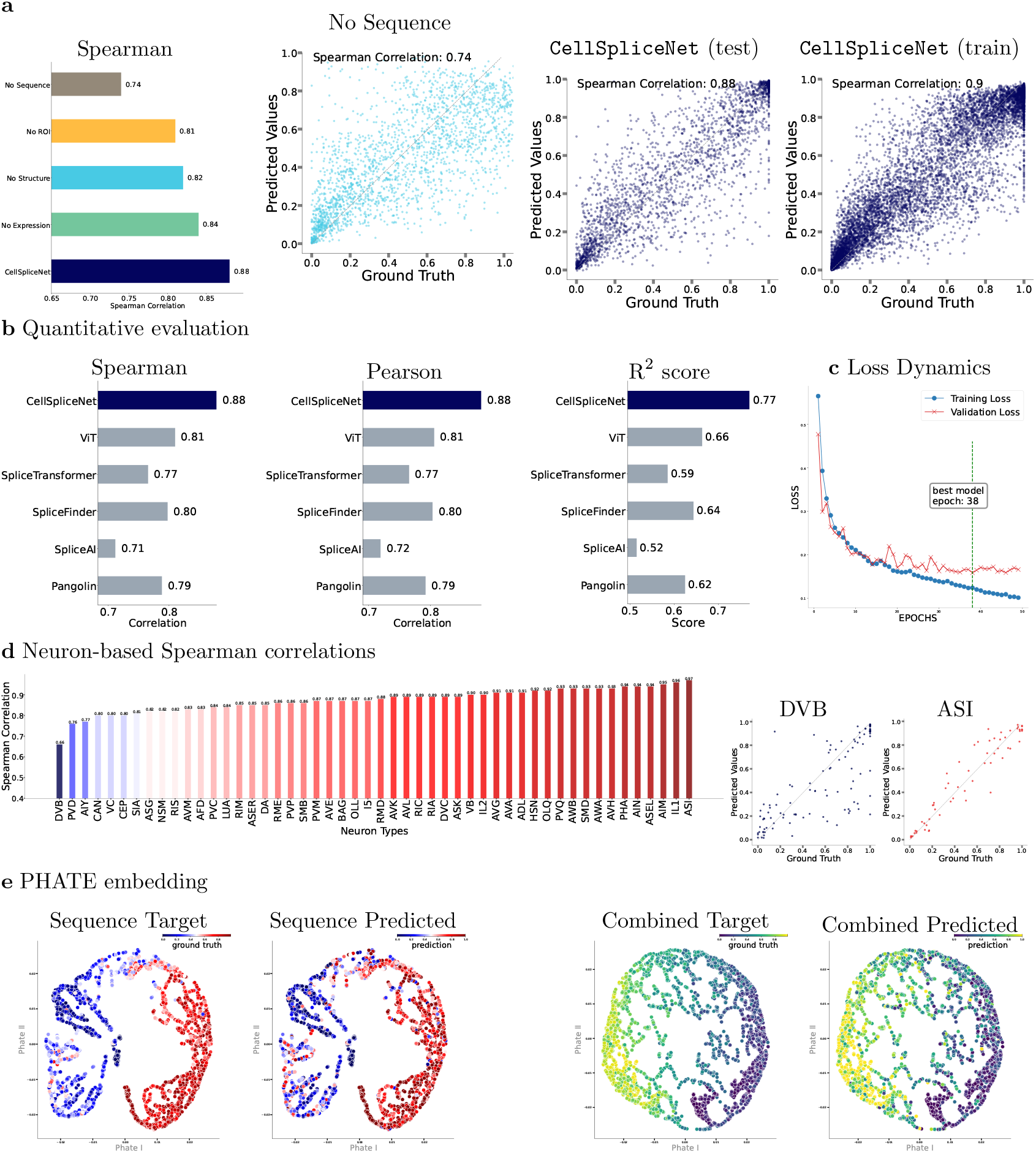
CellSpliceNet outperforms across different conditions and methods. **a**. Ablation study of `CellSpliceNet` on the test set, showing significant performance decline when the RNA sequence modality is excluded, with the highest Spearman correlation of 0.88 observed when all modalities are utilized. The scatter plots illustrate the impact of excluding modalities in `CellSpliceNet`. Each point corresponds to a test sample. **b**. Comparative analysis of `CellSpliceNet` against five advanced models using R^2^ score, Pearson, and Spearman correlations. **c**. Training and validation loss trends during `CellSpliceNet`‘s development. **d**. The Spearman correlation for `CellSpliceNet` varies across neuron types, and scatter plots display events for the neurons with the lowest (DVB) and highest (ASI) Spearman correlations. **e**. PHATE visualizations of test samples with sequence and combined modalities, color-coded by prediction accuracy and deviations from true values.

#### CellSpliceNet achieves state-of-the-art performance in splicing prediction

To benchmark the performance of `CellSpliceNet` relative to leading approaches, five widely recognized splicing prediction models (ViT [30], SpliceTransformer [29], SpliceFinder [28], SpliceAI [27], and Pangolin [35]) were evaluated using three standard performance metrics: Spearman correlation, Pearson correlation, and coefficient of determination (R^2^). As shown in Fig. 2b, `CellSpliceNet` consistently outperforms all baselines. It leads the general-purpose ViT model by 0.07 in both correlation metrics and by 0.11 in R^2^. Compared to the four domain-specific models, the margin of improvement is even larger. For fair comparison, all models were trained using three independent random seeds, and the mean values across repetitions are reported. These findings confirm that the integration of multiple data modalities in `CellSpliceNet` yields significant gains over current state-of-the-art methods.

In addition to overall performance, convergence behavior was examined to assess training dynamics and generalization stability (Fig. 2c). Plots of training (blue) and validation (red) losses over time show a smooth and consistent decline, indicating efficient learning and generalization on held-out data.

To evaluate performance across biologically distinct contexts, the test set was stratified by neuron type, with subtypes ranked by Spearman correlation (Fig. 2d). Although DVB (0.66), PVD (0.76), and AIY (0.77) exhibit relatively lower correlations, all other neuron types exceed 0.80. Particularly high correlations were observed for ASI (0.97), IL1 (0.96), and AIM (0.95). Representative scatter plots for DVB and ASI, respectively corresponding to the lowest and highest correlations, further illustrate `CellSpliceNet`‘s capacity to generalize across subpopulations with diverse transcriptomic profiles.

Finally, to visualize how `CellSpliceNet` organizes PSI-related representations, PHATE [36] embeddings were generated from the test set sequence modality prior to the final MLP layer (Fig. 2**e**). Embeddings are colored by target and predicted PSI values, revealing a close alignment between the learned latent space and the ground truth. Taken together, these results demonstrate that the multimodal architecture of `CellSpliceNet` not only advances the state of the art but also ensures reliable prediction across heterogeneous neuronal cell types.

### Uncovering attention in full-length RNA sequences

To gain insight into which regions of the input are most critical for splicing prediction, we interrogated the attention pooling layer following multi-head attention modules at the core of `CellSpliceNet`. By examining this layer, where each attention head aggregates contextual cues before contributing to the splicing (PSI) estimate, we can visualize how the model distributes its focus across the RNA. This approach highlights specific patches within each modality, in this case, the whole RNA sequence, that exert the greatest influence on splicing decisions. `CellSpliceNet`‘s inferred attention patterns reveal a pronounced focus on intron-exon boundaries across all neuron types examined (Fig. 3a), underscoring its ability to internalize canonical splicing signals. When these attention weights are aggregated around key regions, the histograms in Fig. 3b confirm that the vast majority of attention falls within exonic and immediately flanking intronic patches, in line with well-established roles of splice sites and nearby auxiliary elements in guiding splicing decisions.

**Fig. 3:**
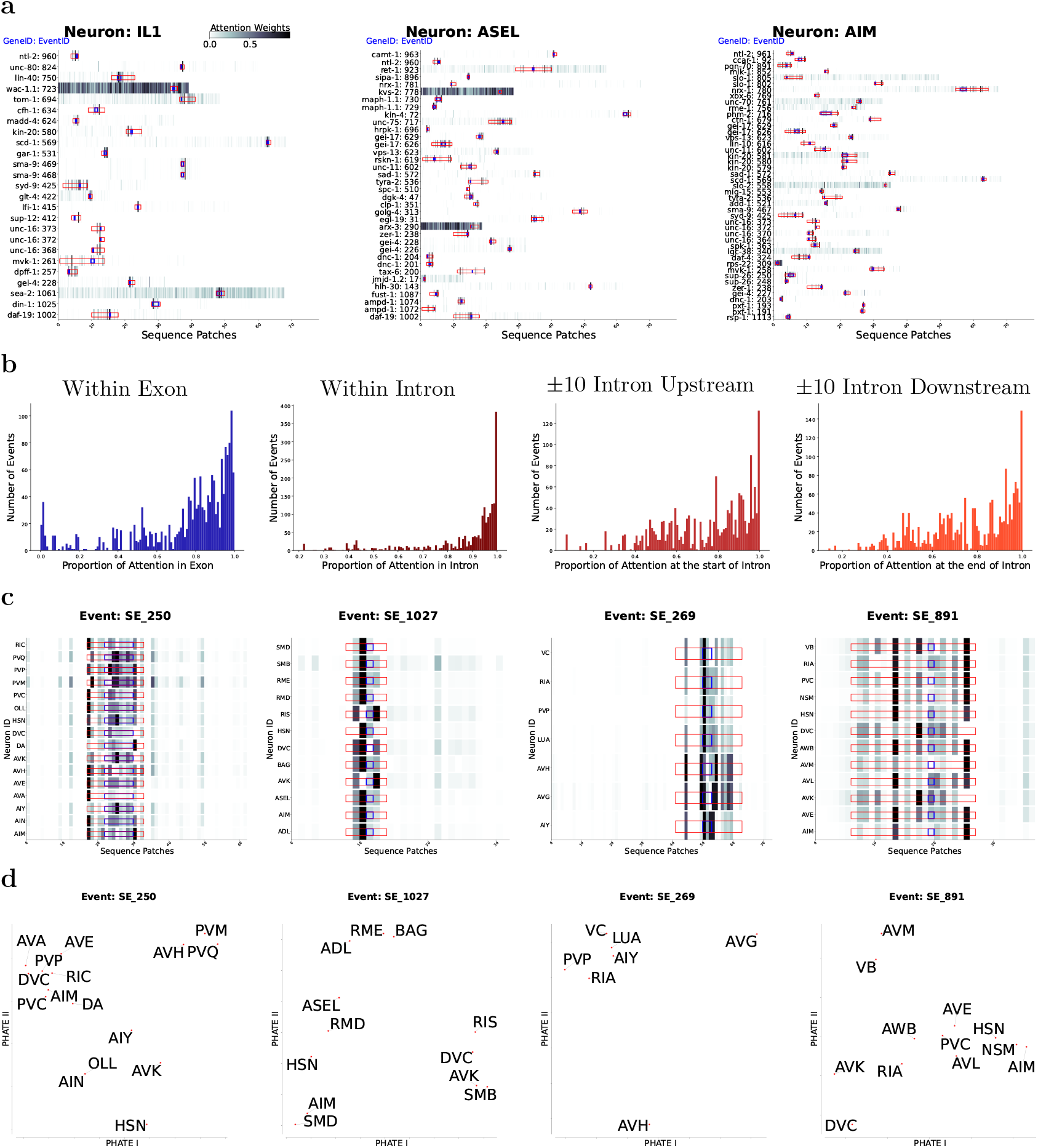
CellSpliceNet infers high attention weights around the exons and introns in RNA Sequence. **a**. Heatmaps (white→black) of attention weights for multiple splicing events across three neuron types, with introns (red) and exons (blue) outlined; `CellSpliceNet` predominantly targets intron-exon boundaries, although some events (e.g., event wac-1.1 in IL1) show a broader sequence focus. Here, a sequence patch is defined as a contiguous, fixed-length segment of base pairs taken sequentially, enabling the model to break long RNA sequences into units while preserving global context. **b**. Histograms of attention fractions falling inside exons, inside introns, and within *±*5 patches upstream or downstream of intron-exon junctions. For every splicing event, we quantified the normalized fraction of model attention assigned to exonic versus intronic regions and visualized the resulting distribution as histograms. **c**. Per-patch attention profiles for four exemplar events across neuron types, revealing both neuron-specific and shared patterns. **d**. PHATE embeddings of these attention signatures cluster neurons with similar profiles, corroborating the observed similarities.

Beyond these universal trends, Fig. 3c presents four representative splicing events and contrasts per-patch attention profiles across neuron types: while certain neurons (e.g., X and Y) exhibit remarkably similar distributions of attention weights, others display distinct, event-specific patterns that likely reflect divergent regulatory contexts. This variability suggests that `CellSpliceNet` not only memorizes conserved boundary features but also dynamically reallocates attention to capture subtle, neuron-dependent cues.

Fig. 3d reinforces these cross-neuron differences by embedding each neuron’s attention signature into a low-dimensional PHATE space. Neurons with analogous attention profiles cluster together for each event, visually corroborating the patterns seen in Fig. 3c. Taken together, these results demonstrate that `CellSpliceNet` strikes a balance between recognizing universal splicing motifs and adapting its focus to the unique transcriptomic landscapes of different neuron types.

Four histograms quantify where the model allocates its attention: 1) proportion of total attention that falls inside exons, 2) attention proportion to intronic sequence, 3) attention weight proportion at ± 5 patches upstream of the intron, and 4) attention concentrated at the downstream of the intron (± 5 patches). (c) Cross-neuron comparisons for four representative events. Each subplot lists neuron types on the y-axis and shows per-patch attention bars predicted by `CellSpliceNet`. While the model often tailors its focus to neuron-specific splicing patterns, certain neurons exhibit strikingly similar profiles. The PHATE embeddings plotted directly beneath (Fig. 3d) visually corroborate these similarities by clustering neurons with analogous attention signatures for each event.

### Interpreting attention in RNA structure

The structural modality in `CellSpliceNet` is centred on the midpoint of each exon and extends symmetrically into the flanking intronic regions, providing both nucleotide-level and secondary-structure context. We therefore dissected its attention landscape along two complementary axes: primary sequence position and base-pairing topology.

#### Positional profile

Across representative neurons (OLQ, AFD, PVQ, and DA), attention densities converge sharply on the exon core (Fig. 4a). Salience diminishes monotonically with distance from the splice sites, indicating that structural cues within the exon itself dominate the model’s PSI estimates, whereas more distal intronic structures are comparatively uninformative.

**Fig. 4:**
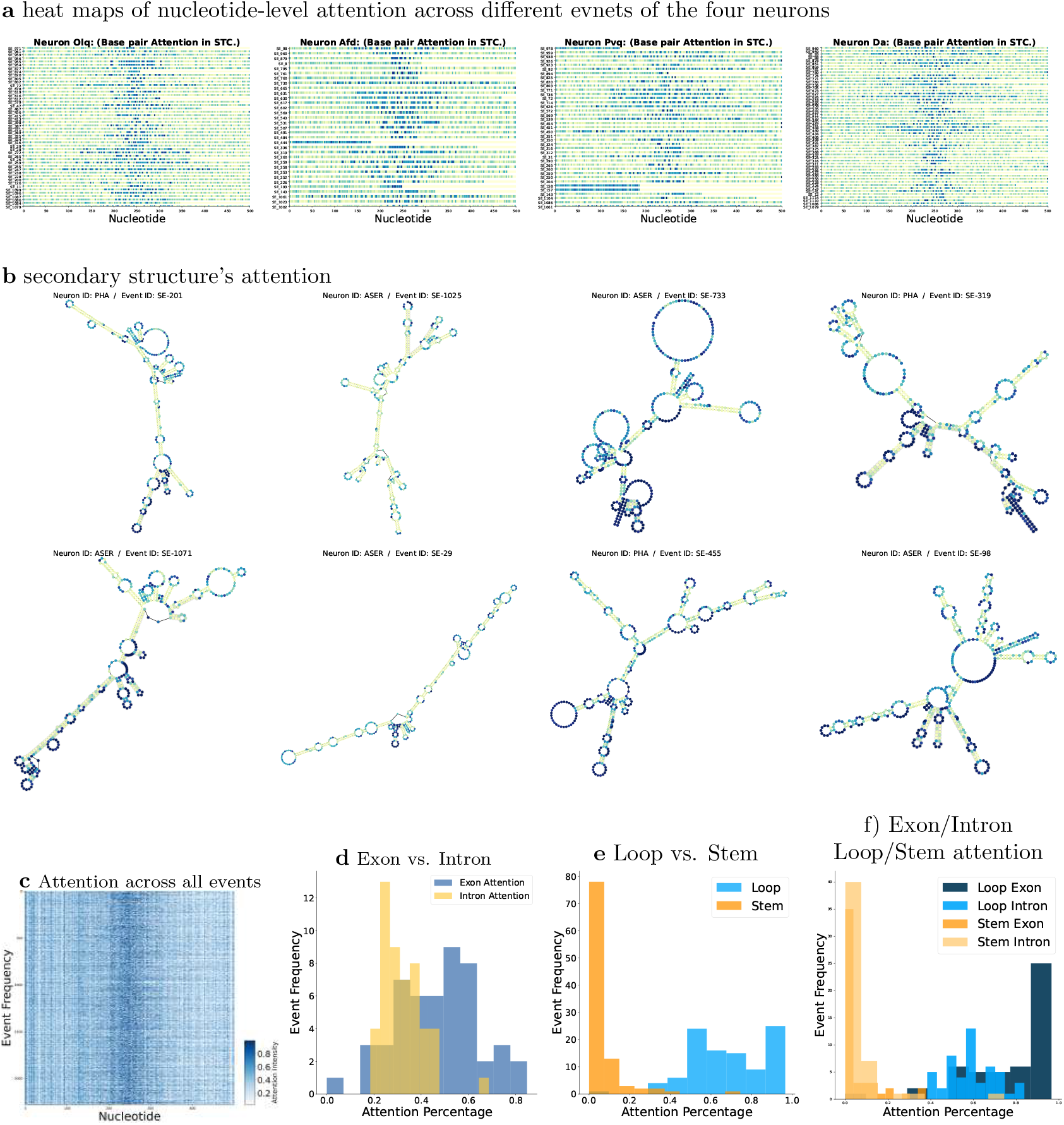
Structure-modality attention concentrates on exonic loop nucleotides. **a**. Heat maps of nucleotide-level attention for four exemplar neurons (OLQ, AFD, PVQ, DA) reveal a sharp focus on the central cassette exon, with salience tapering off into the flanking introns. **b**. Event-based attention over the secondary-structure overlays for two representative events shows that peaks (blue) coincide with unpaired loop bases, while stem regions are down-weighted. **c-d**. Meta-profile frequencies obtained by stacking all annotated events across the nervous system confirm a genome-wide maximum at the exonic core. **e-f**. Histograms comparing attention weights assigned to loop versus stem nucleotides demonstrate a pronounced enrichment on loops, especially within exons, consistent with a model in which accessible single-stranded motifs drive alternative splicing decisions.

#### Topology-specific preferences

Fine-grained mapping of attention onto predicted secondary structures (Fig. 4b) reveals a pronounced bias towards unpaired nucleotides. Loop regions (highlighted, for example, in ASER (event SE-1025) and PHA (event SE-201)) exhibit punctate peaks, while adjacent stems are systematically de-emphasized. This pattern holds across the genome: when all neuronal events are aggregated, attention reaches its global maximum within loop nucleotides located in the exonic core (Fig. 4c-d). Quantitative comparison confirms that loop residues receive substantially higher weights than stem residues, with exonic loops outscoring stemmed pairs by a wide margin (Fig. 4e-f).

Collectively, these observations support a model in which accessible, single-stranded motifs inside the exon provide the principal structural signals guiding alternative splicing decisions, whereas double-stranded stems contribute only marginally. The structural modality of `CellSpliceNet` thus recapitulates and extends classic biochemical insights into the role of RNA conformation in splice-site selection.

### Analyzing nucleotide-level and motif-level attention in the regions of interest

The ROI stream in `CellSpliceNet` zooms in on the two splice junctions that flank every cassette exon: the upstream intron-exon boundary and the downstream exon-intron boundary. Because splicing fidelity hinges on short sequence motifs located within these narrow windows, the ROI attention-pooling layer offers a uniquely fine-grained view of how the network weighs individual nucleotides and their associated regulatory elements.

#### Local inspection in representative neurons

We first visualised nucleotide-level attention for four prototypical neurons (Fig. 5a; additional examples in Supplementary). Across all cases, attention peaks sharply over the exon core and tapers into the flanking introns, mirroring the distribution of exonic splicing enhancers and the rapid decay of regulatory signal with distance from the splice site. Notably, the upstream region attracts stronger weights than its downstream counterpart, hinting at a model preference for features that distinguish exon definition at transcript entry rather than exit.

**Fig. 5:**
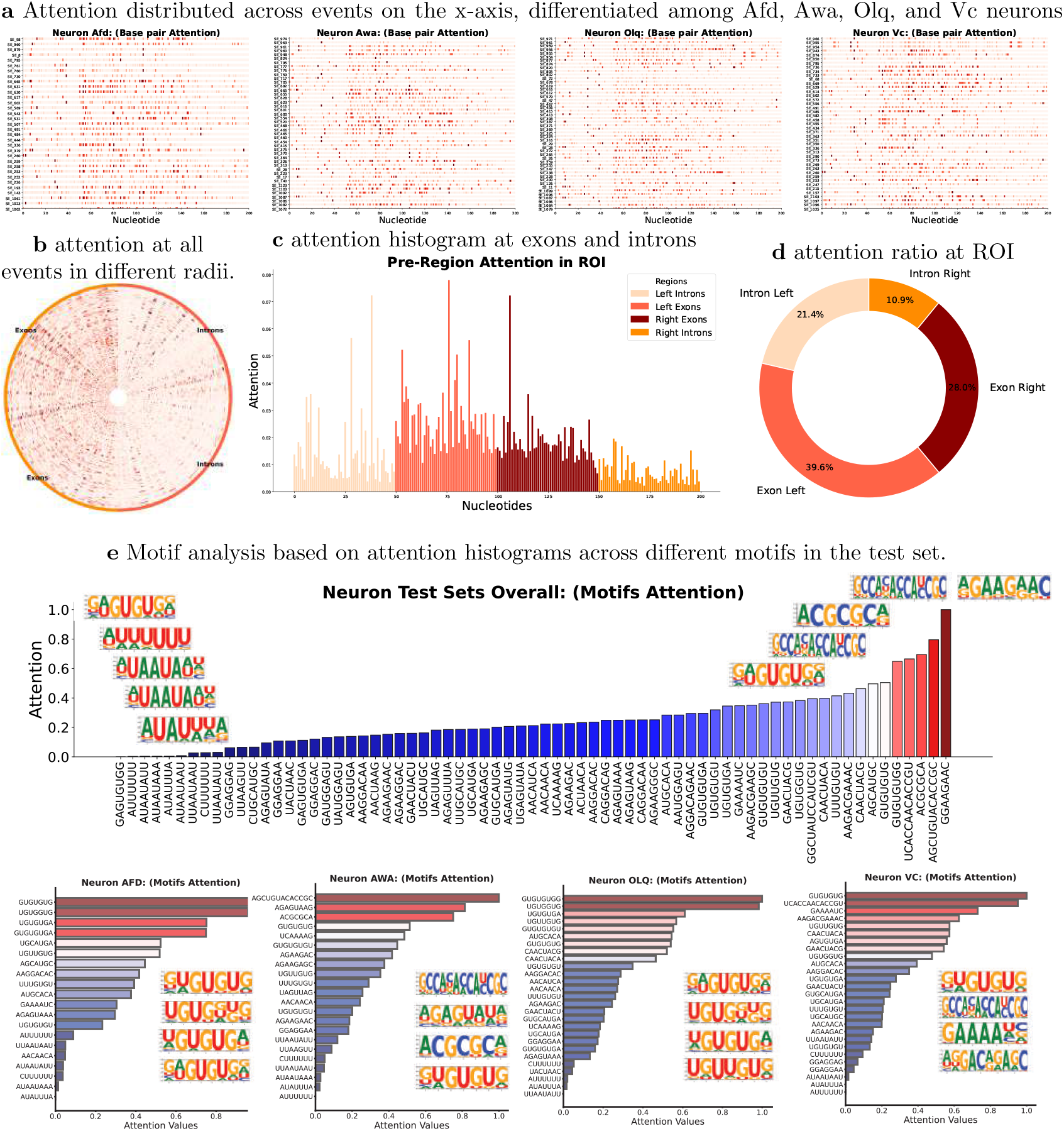
Elevated attention at upstream intron-exon boundary and GGAAGAAC motif. **a**. displays nucleotide-level attention across representative neurons, highlighting a preference for upstream exon regions. **b**. Attention distribution as an “attention wheel” shows an upstream bias. **c**. Quantification of this asymmetry, separating attention into different genomic compartments. **d**. Aggregated attention data shows dominant upstream exonic signals. **e**. Sequence motifs are rankedby their attention weight, revealing key enhancers. **f**. Stratified motif attention by neuron class suggests neuron-specific splicing codes.

#### Global landscape across the test set

To summarise patterns over the entire held-out cohort, we arranged every cassette event on a concentric “attention wheel”, assigning each exon a unique radius and mapping attention as angular intensity (Fig. 5b). A clear dichotomy emerges: the left hemispheres glow more intensely than the right hemispheres, underscoring a systematic upstream bias. This asymmetry is quantified in Fig. 5c, which separates attention values into four compartments—left exon, left intron, right exon, right intron—and displays their distributions as density plots.

#### Quantitative enrichment of upstream exonic signal

Aggregating over all test events (Fig. 5d), the upstream exon sequesters 39.6% of the total ROI attention, followed by the downstream exon (28.0%), the upstream intron (21.4%), and, trailing far behind, the downstream intron (10.9%). That the left exon outcompetes every other compartment is striking, given the model’s symmetry in architecture and training objective. One plausible explanation is that upstream exon definition in *C. elegans* relies heavily on strong splicing enhancers and the consensus GU dinucleotide at the donor site, whereas the acceptor AG and associated polypyrimidine tracts, though essential, are more stereotyped and therefore less informative for discrimination. Consistent with this view, motif analysis of the top-weighted nucleotides reveals recurrent enrichment for canonical ESE hexamers (e.g. `GAAGAA`, `GACGGT`) in the left exon, corroborating experimental reports that upstream enhancers modulate exon inclusion probability.

#### Functional implications

Together, these observations indicate that `CellSpliceNet` internalises a mechanistic hierarchy reminiscent of classical splice-site biology: it first anchors on the upstream exon to gauge splicing competence, then refines its estimate through diminishing contributions from the downstream context. By highlighting the exact nucleotides and motifs that carry predictive weight, the ROI attention lens not only enriches our understanding of alternative splicing logic but also nominates specific sequence features for future experimental validation.

#### Motif-level resolution of ROI attention

Beyond the base-wise view, we collapsed the ROI attention map onto all 8-14 nt sequence motifs present in the cassette events and ranked them by cumulative weight. The resulting distribution (Fig. 5e) is highly skewed: the enhancer-like octamer GGAAGAAC accounts for the single largest share of attention, followed by AGCUGUACACCGC, `ACGCGCA`, UCACCAACACCGU, and GUGUGUGG. All five harbour purine-rich cores or UG dinucleotide repeats that have previously been implicated in splicing activation or repression, underscoring the biological plausibility of the model’s focus.

When motif weights are stratified by neuron class (Fig. 5f), distinct regulatory preferences surface. For instance, the UG-repeat heptamer `GUGUGUG` dominates AFD and VC neurons, whereas AGCUGUACACCGC is most prominent in AWA. Likewise, the octamer GUGUGUGG emerges as the top-ranked motif in QLQ. These shifts suggest that neuron-specific splicing codes may be driven by differential deployment of the same repertoire of regulatory words. Comprehensive motif-to-neuron mappings are provided in Supplementary Table 3, offering a resource for targeted perturbation experiments aimed at deciphering cell-type-specific splicing logic.

### Understanding attention in gene expression through splicing-factor dynamics

We have demonstrated how `CellSpliceNet` captures the contributions of whole RNA sequence, local structure, and specific regions of interest. We now turn to the expression modality, interrogating the attention weights assigned to individual splicing factors across the *C. elegans* distinct neurons. By aggregating these attentional scores over neuron classes, the model pinpoints the regulators that are most predictive of the PSI metric, clarifying the extent to which each neuron type depends on distinct components of the splicing apparatus.

`CellSpliceNet` reveals marked heterogeneity in splicing-factor usage (Fig. 6a). Some neurons enlist a broad repertoire of regulators, whereas others rely on a narrowly defined subset. Hierarchical clustering of factors whose attention values fall within the top 85th percentile (Fig. 6b) crystallizes this contrast. Highly influential factors, namely *smu-1, unc-75, hrp-1, exc-7, grld-1, snr-4, fox-1, rnp-3, pes-4*, and *rsp-6*, dominate the regulatory landscape of many neurons, whereas *pab-2, hrp-2, prp-6, nova-1, fust-1, fubl-1, rbm-25, asd-2, pab-1*, and *rsp-1* are consistently down-weighted.

**Fig. 6:**
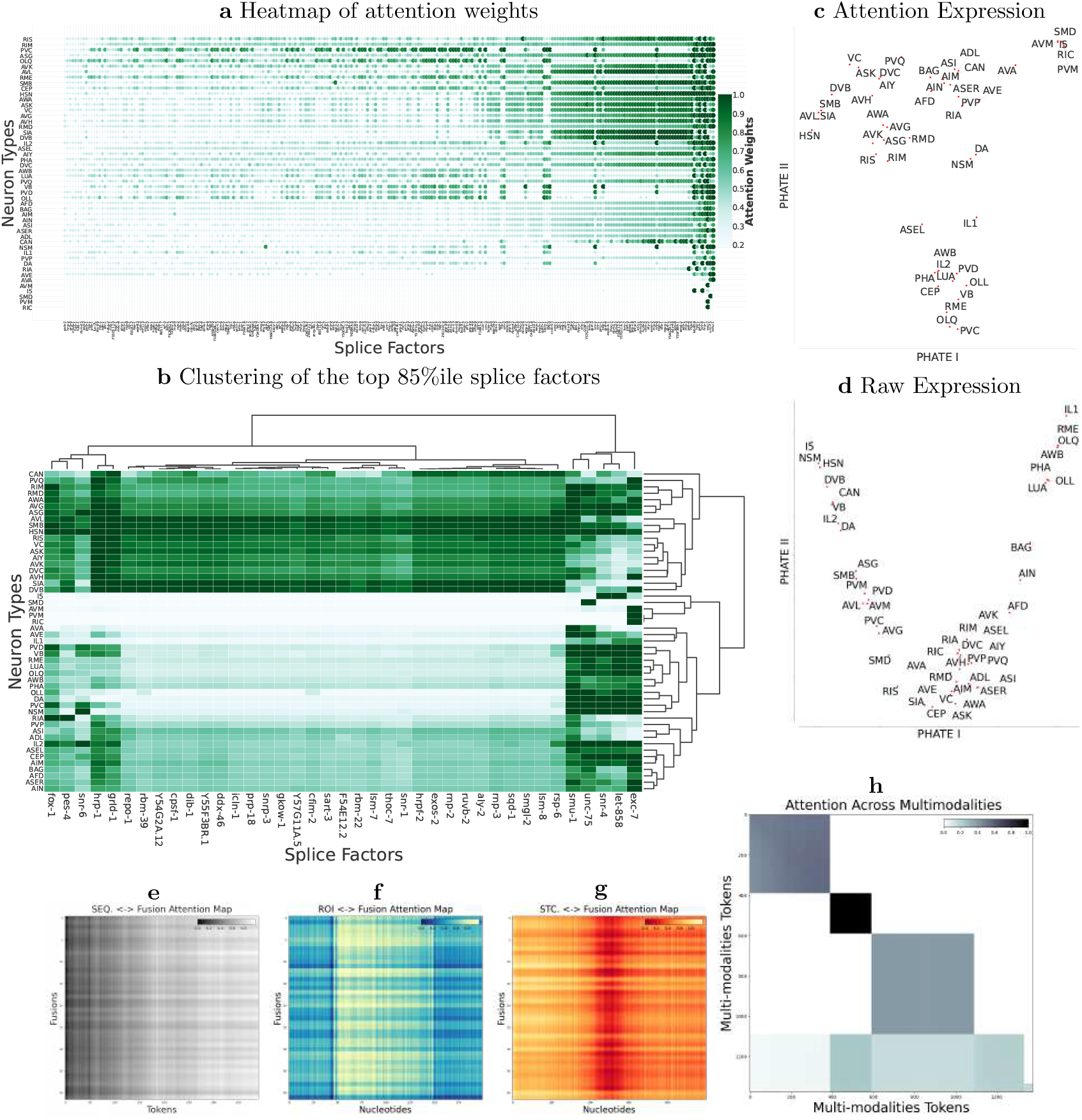
Gene Expression and Fusion Attentions Inference by CellSpliceNet. **a**. Presentation of inferred attention weights for splicing factors across different neuron types, arranged in columns to depict the distribution and intensity of attention. The splicing factors are organized in descending order of impact, from the highest to the lowest, from the first row to the last. **b**. Hierarchical clustering of splice factors, focusing on the top 85th percentile to reveal patterns and relationships based on expression levels and interactions. **c-d**. PHATE embedding visualization of gene expression profiles across different neuron types in attention expression and raw expression, respectively, highlighting the multidimensional expression landscape and facilitating the identification of distinct expression patterns and clusters. **e-g**. Attention maps for the fusion modality of `CellSpliceNet`. The model assigns its highest weights to the sequence termini, down-weights exon-intron junctions, and largely ignores central exons, whereas the expression channel distributes attention uniformly. **h**. Mean attention across the entire test set, summarizing modality-specific attention patterns.

From the neuron’s perspective, neuron classes such as CAN, AWA, AVG, ASG, PVQ, RIM, RMD, AVL, SMB, and DVC display diverse portfolios of highly weighted factors, indicative of multiple convergent regulatory routes. Conversely, AVM, PVM, RIC, I5, SMD, AVA, AVE, IL1, and OLL focus their regulatory attention on comparatively few factors, suggesting streamlined splicing control. These patterns are immediately apparent in the heat map, where rows of splicing factors ordered by attention values expose neuron-specific signatures.

Projecting the splice factors profiles into a PHATE manifold (Fig. 6c) resolves three discernible clusters of neuron types, each apparently governed by a shared regulatory programme. The upper-left cluster comprises I5, NSM, HSN, DVB, CAN, VB, IL2, and DA, reflecting a cohesive splicing strategy; a central cluster assembles RMD, AIM, ADL, RIC, DVC, and PVP; and a far-right cluster includes IL1, RME, OLQ, AWB, PHA, OLL, and LUA. This geometry underscores lineage- or function-specific splicing regimes and highlights the power of attention-based interpretability to map regulatory logic across an entire neuronal repertoire.

### Interpreting cross-modal attention in the fusion

Multi-modal integration in `CellSpliceNet` is mediated by a dedicated fusion modality - a trainable query bank that interrogates every other modality and reconciles their signals. Analysis of its mean attention weights (Fig. 6d) uncovers a distinctive and highly unique pattern. Overall, the attention weights for the ROI and expression channels are higher compared to those of the whole sequence and structure channels. Additionally, within the full-sequence channel, fusion queries progressively increase their weight from the beginning of the sequence toward the end of it, in sharp contrast to the uniform positional emphasis observed in the sequence encoder itself. In the region-of-interest channel, they deliberately down-weight canonical exon-intron and intron-exon junctions, and in the structural channel, they eschew the central exon, instead sampling flanking regions. By comparison, attention in the expression channel is distributed almost uniformly.

Strikingly, these choices run differently from the priorities learned and discussed within each modality in isolation, implying a purposeful division of labour. We propose that the modality-specific encoders forward their most decisive features directly to the prediction head, whereas the fusion layer seeks complementary, cross-modal cues that resolve residual uncertainty. In this view, the fusion modality performs a fine-grained search for corroborative signals that lie outside the high-salience regions already captured by the individual encoders, thereby sharpening the model’s final PSI estimates.

## Discussion

We introduce `CellSpliceNet`, a multimodal transformer framework that unifies full-length sequence, secondary structure, splice-junction context, and single-cell expression to predict alternative splicing with unprecedented accuracy and interpretability. By jointly embedding these complementary views of an RNA molecule and its regulatory milieu, `CellSpliceNet` achieves a Spearman correlation of 0.88 on held-out samples—a significant gain over the strongest single-modality models and a clear margin over five state-of-the-art baselines. Equally important, the network’s attention landscapes recapitulate—and extend—decades of biochemical insight, pinpointing canonical boundary motifs, exon-centric loop residues, and enhancer-rich upstream nucleotides as dominant drivers of exon inclusion. These results highlight the power of cross-modal fusion for decoding complex post-transcriptional regulation. Building on the Zorro framework, we refine the multimodal multihead-attention blocks so that each modality first learns “pure” internal representations while a bank of learnable fusion queries coordinates cross-talk. Crucially, our masking scheme grants the expression stream privileged connectivity: splicing-factor tokens can broadcast cell-wide regulatory signals directly to sequence, structure, and ROI encoders, and again through the fusion layer. This targeted relaxation of modality isolation allows `CellSpliceNet` to inject neuron-specific context precisely where it matters, elevating performance without diluting the distinct inductive biases of the other modalities. After encoding, modality-specific attention-pooling networks compress hundreds of tokens into compact, weighted embeddings. The learned coefficients illuminate the regulatory hierarchy: whole-sequence and ROI pools concentrate weight on intron–exon boundaries; structure pools privilege single-stranded loops inside exons; and expression pools highlight a small cadre of high-impact splice factors. These pools are not mere conveniences for dimensionality reduction; ablation shows that eliminating any one of them, or replacing them with naïive mean pooling, erodes both prediction accuracy and interpretability. The model’s attribution maps reinforce classical splicing dogma while exposing subtle, cell-type-specific nuances. Its upstream exon bias, structural loop preference, and dynamic boundary focus all arise spontaneously from the end-to-end loss—strong evidence that the masking-plus-pooling strategy guides the network toward biologically grounded solutions. Moreover, allowing expression tokens privileged access appears vital: when their direct connections are ablated, downstream attention to enhancer-rich nucleotides wanes and predictive power falls, underscoring the global influence of splice-factor abundance.

The interpretable embeddings and pooling weights produced by `CellSpliceNet` offer a direct path to prioritising disease-associated variants, designing antisense oligonucleotides, and charting cell-type-specific splicing networks. Because the masking architecture is generic, it can be transplanted to other multimodal genomics problems—such as integrating sequence, epigenome, and 3D-contact maps for transcriptional prediction—where preserving modality purity while enabling selective cross-talk is equally desirable.

`CellSpliceNet` demonstrates that precise masking, modality-specific pooling, and biologically informed loss functions can transform raw multimodal data into accurate, interpretable models of RNA regulation. By revealing how global splice-factor context shapes local sequence decisions, the framework lays the groundwork for a new generation of integrative, mechanism-aware predictors in molecular biology.

## Methods

### Overview

Building on the need to model alternative splicing with multimodal data, we detail below the data preprocessing steps, embedding pipelines for each modality, multimodal transformer architecture, pooling and prediction mechanisms, and training procedures. This section aims for reproducibility, while background motivations and rationales are provided where necessary.

### Data sources and preprocessing

*C. elegans* possesses a remarkably simple yet well-characterized nervous system, consisting of 302 neurons partitioned into 118 distinct anatomical classes. White et al. [37] achieved the first complete mapping of these neurons and their connections, providing a foundational “connectome” that remains unique in its cellular resolution. This comprehensive wiring diagram, together with the worm’s invariant developmental lineage and genetic tractability, makes *C. elegans* uniquely suited for systems-level neuroscience. In particular, recent technical advances have yielded broad and well-annotated neuron-specific RNA sequencing datasets,effectively a gene expression atlas spanning the entire nervous system [38]. Such resources now enable investigators to profile every neuron’s transcriptome and interrogate complex regulatory phenomena (for example, neuron-specific alternative splicing) with single-cell precision [39].

In this work, we employed the high-quality gene and transcript annotations provided by Wormbase [40]. For each target exon, intronic regions upstream and downstream were extracted to analyze local sequence context and predict RNA secondary structure. Secondary structures were computed using ViennaRNA [32] with default parameters. Additionally, we utilized transcriptomic data generated by the CeNGEN Consortium[41, 42], which contains RNA sequencing data providing full gene coverage across 50 types of isolated neurons, to quantify cell-type-specific exon usage[39], complemented with single-cell sequencing data[38] to quantify gene expression. Raw count matrices were preprocessed by quality control filtering (e.g., minimum gene/cell detection thresholds), normalization (e.g., library-size scaling), and imputation using MAGIC [17] to mitigate dropout effects. A curated list of 243 splice factors was compiled based on prior literature and databases.

### Modality-specific embedding in CellSpliceNet

We designed four parallel embedding pipelines for sequence, region-of-interest (ROI), secondary structure, and gene expression modalities. The embedding pipelines produce fixed-dimensional token embeddings for integration in the multimodal transformer.

#### Sequence embedding

The advent of genomic data availability due to advances in sequencing technologies in the last two decades positions it as an ideal candidate for deep learning applications. However, several challenges have historically impeded progress in this domain. A primary issue is the need to encode exceedingly long sequences to capture essential long-range biological interactions [43]. The computational demand increases significantly for transformers because their attention mechanisms scale quadratically with sequence length [8]. Recently, K-mer tokenization strategies have been employed to address this computational challenge [44, 45, 46]. Additionally, due to the biological significance of single nucleotide phenomena (e.g., SNPs, splice sites), there is a growing interest in models that consider individual nucleotides as tokens. Various models, including transformers [47] and other architectures [48, 26], have incorporated a data preprocessing step that truncates sequences to a manageable few hundred nucleotides. Notably, Nguyen et al. [49] implemented a scalable transformer architecture that accommodates sequence lengths up to 1M nucleotides using a more efficient convolution-based mechanism, replacing the traditional attention architecture [50].

We divide the input RNA sequence into *N* patches 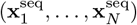, each of a defined `window _size`. If the last patch is shorter than the `window_size`, it is padded for uniformity. Specifically, each patch 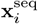 contains the base pair’s type and annotation tokens (e.g., exon, intron, or flanking regions), which are one-hot encoded and concatenated with the encoded RNA base pairs.

To embed each patch, 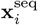 undergoes a transformation via a linear layer with parameters 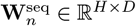 and 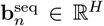, followed by layer normalization (LayerNorm) and the addition of a learnable positional embedding **E**^seq^, described in Eqn ().

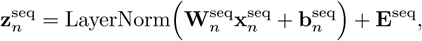

where 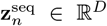. This methodology effectively retains critical sequence and annotation information, facilitating the extraction of meaningful insights by downstream transformer blacks of our `CellSpliceNet`.

#### Structure embedding

RNA secondary structure influences splicing by modulating the accessibility of key regions such as splice sites and regulatory motifs. The embedment of these regions in stable structures, such as hairpins or stems, reduces their accessibility to the splicing machinery, potentially altering splicing patterns and generating diverse transcripts from the same RNA sequence.

We model RNA secondary structure over a window size of *S*, where the window’s middle is at the center of exon *f*, focusing exclusively on essential structural features around splicing regions, particularly the flanking introns and the encompassed exon. We utilize ViennaRNA [32] to fold these sequences into two-dimensional secondary structures, which are then stored as graphs. In a similar approach to our gene expression graphs, we apply a geometric scattering transform as described below, with Dirac signals on these graphs, encoding multiscale RNA secondary structures through cascading graph wavelets.

To embed the secondary structure modality, let 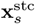 denote the raw scattering coefficients at structure position *S*. These coefficients undergo a linear transformation followed by normalization and positional encoding. Specifically, the coefficients are first transformed by a linear layer with learnable parameters 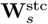 and 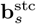. The output is then normalized using Layer Normalization and enhanced with a learnable positional embedding **E**^stc^, as illustrated in Eqn (2).

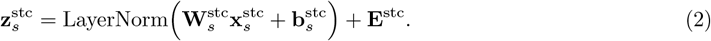

This process yields the final embedded representation 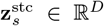, encapsulating both the transformed scattering coefficients and their contextual position within the structural modality.

#### ROI embedding

While tokenizing long RNA sequences mitigates the computational challenges [45, 46] faced by the `model` in processing extensive sequences, analyzing self-attentions across base pairs poses difficulties for downstream analysis and motif interpretation within the region of interest (ROI). In our approach, we first select *B* number of base pairs (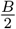 from the beginning and 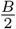 from the end of exons) and represent the discretized base pairs at position *j* within the ROI as 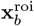. Each base pair position undergoes a dedicated linear transformation followed by layer normalization to compute its embedding. Specifically, the embedding for each position is given by Eqn ().

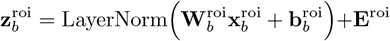

Here, 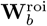 and 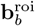 represent the weight matrix and bias vector of the linear layer for the b-th base pair in the ROI, respectively. Additionally, 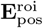 from ℝ^*H*^ provides positional context for the j-th base pair. This process yields an embedding 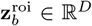, encapsulating the critical attributes of base pairs at each position and furnishing a detailed representation of the ROI within the defined high-dimensional space.

#### Gene expression embedding

To characterise pairwise relationships among splice factors in single-cell transcriptomic data we employ a multi-step pre-processing and graph-based analysis pipeline. Let **E** ∈ ℝ^*F* ×*C*^ be the raw count matrix, with *F* splice factors measured across *C* cells. We first impute missing entries with Markov Affinity-based Graph Imputation of Cells (MAGIC) [17], obtaining an enhanced gene expression matrix **E**_MAGIC_. Retaining only splice factors relevant to our study yields **E**_masked_ ∈ ℝ^*f* ×*C*^ with *f* ≤ *F*.

The masked matrix is partitioned into *K* neuron-type subsets 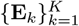, where 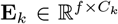. For each type we calculate pairwise mutual information (MI) between splice factors, forming a symmetric matrix **M**_*k*_ ∈ ℝ^*f* ×*f*^; entry [**M**_*k*_]_*ij*_ quantifies MI between factors *i* and *j*. We treat **M**_*k*_ as the weighted adjacency of a splice-factor co-expression graph and assign each node a scalar signal equal to the mean expression of that factor across cells, **a**_*k*_ ∈ ℝ^*f*^.

Applying the geometric scattering transform (see supplementary matrial and Fig. 1 for a schematic of the geometric embedding of the expression modality) to each MI-weighted graph produces a multiscale coefficient tensor **C**_*k*_ ∈ ℝ^*f* ×*S*×*d*^. We use Dirac signals weighted by average gene expression as the node features for this process. Passing **C**_*k*_ through a multi-layer perceptron (MLP) yields splice-factor embeddings **Z**_*k*_ = MLP(**C**_*k*_); stacking across neuron types gives **Z** ∈ ℝ^K×hidden dim^.

To obtain a single expression vector per neuron type we use an attention mechanism. Let **z**_*k*,*i*_ ∈ ℝ^*D*^ be the embedding of splice factor *i* for type *k* and let ***α*** ∈ ℝ^*K*×*S*^ be a learnable parameter matrix. Row ***α***_*k*_ is normalised with a softmax to give weights **a**_*k*_ = softmax(***α***_*k*_). The neuron-type expression embedding is then

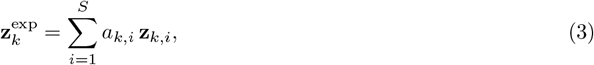

which highlights the most informative splice factors for type *k*.

### Multimodal transformer and Zorro attention

`CellSpliceNet` uses a multimodal multi-head transformer to integrate the four aforementioned modalities along with a set of learnable fusion tokens. While conceptually related to a recent work on multimodal protein analysis [51], our approach introduces a more advanced techniques for information fusion.

As previously described, the total number of tokens for the {sequence, structure, ROI, expression} modalities are {*N*, *S, B, S*} respectively, where each token is embedded into ℝ^*D*^. Concretely, we denote:

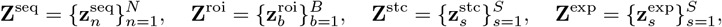

where **z** ∈ ℝ^*D*^. In addition, we introduce *P* learnable fusion tokens: 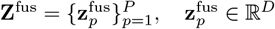.

The multimodal tokens are formed by concatenating the modality-specific tokens from each individual modality **Z**^multimodal^ = **Z**^seq^||**Z**^stc^||**Z**^roi^||**Z**^exp^||**Z**^fus^ ∈ ℝ^*(N +2S+B+P)*×*D*^.

In a vanilla transformer [8], a standard multi-head self-attention module will be used to compute key **K**, query **Q**, and value **V** as learned projections of **Z**^multimodal^. However, this assumes universal and bidirectional attention, which does not accurately model the correct information exchange over the five modalities. To do this, we used a `Zorro` attention mechanism [33], which accommodates specific attention constraints.

Specifically, let modality(*i*) ∈ {seq, stc, roi, exp, fus} denote the modality type of the *i*-th token. We define a `Zorro` mask **MASK** ∈ {0, 1}^*L*×*L*^, where *L* = *N* + 2*S* + *B* + *P*, is defined by:

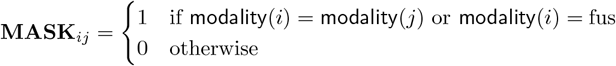

With the `Zorro` mask, fusion tokens attend to all tokens, whereas sequence, structure, ROI, and expression tokens attend only to themselves or fusion tokens. The final multimodal hidden state is produced through the Zorro attention operation over all modalities:

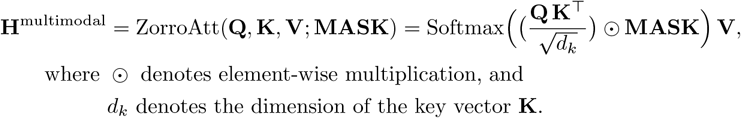

In implementation, we first embed or project raw data from each modality into token form and then concatenate these tokens along with the learnable fusion tokens. We generate the `Zorro` mask by comparing modality types. This masked attention is repeated across multiple heads and stacked for several transformer blocks, leading to final context-rich embeddings from all four modalities, mediated by the fusion tokens.

### Attention-based pooling and PSI prediction

Processing the tokens with the multimodal transformer module is a dimension-preserving step, which means **H**^multimodal^ ∈ ℝ^*(N +2S+B+P)*×*D*^. We can therefore re-partition the multimodal hidden state by modality. This gives us the sequence hidden state (**H**^seq^ ∈ ℝ^*N* ×*D*^), structure hidden state (**H**^stc^ ∈ ℝ^*S*×*D*^), ROI hidden state (**H**^roi^ ∈ ℝ^*B*×*D*^), expression hidden state (**H**^exp^ ∈ ℝ^*S*×*D*^), and fusion hidden state (**H**^fus^ ∈ ℝ^*P* ×*D*^).

For each modality, we apply attention pooling to aggregate the hidden state into a single vector, following RELSO [34]. Concretely, for each modality-specific hidden state **H**^mod^, we learn a low-dimensional attention score:

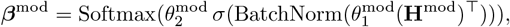

where *β*^mod^ is the attention weights across the input for the given modality, 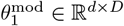 and 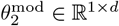 are learned parameters, BatchNorm denotes a batch normalization layer, and *σ*(·) denotes a GELU activation. The resulting attention weights *β*^mod^ are applied to **H**^mod^ to produce a single hidden state vector:

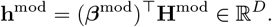

We obtain pooled hidden states for sequence, structure, ROI, expression, and fusion. These are concatenated:

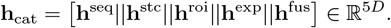

We then feed this concatenated vector **h**_cat_ into a *k*-layer MLP, and the output of the MLP is subsequently passed through a linear layer. The final prediction is obtained using a LogSigmoid activation function, effectively mapping the output to the log-probability space.

This formulation and design helps stabilize training and is particularly beneficial for handling data imbalance and extreme prediction values.

### Training objective

We optimize the model using the loss function defined in Eqn (1), known as the Mean Squared Error (MSE) loss, which penalizes the average squared difference between the estimated values and the actual values.

The choice of MSE as a loss function is motivated by its sensitivity to larger errors, as it squares the differences before averaging them. This property makes MSE particularly useful in scenarios where it is critical to avoid large errors, as the squaring process heavily penalizes larger deviations more than smaller ones, thereby encouraging the model to avoid large errors in its predictions.

### Implementation details

All models are implemented in PyTorch. All experiments are conducted on a single A100 GPU. Data loading and preprocessing pipelines are implemented with standard libraries. Reproducibility is ensured via fixed random seeds and environment specification. Preprocessing scripts, end-to-end training and inference scripts, and pretrained model checkpoints are available in the public repository.

## Data Availability

The datasets from this study can be accessed on the project’s GitHub repository (https://github.com/KrishnaswamyLab/CellSpliceNet-dataset).

## Code Availability

Implementations of the models and optimization algorithms are available at the project’s GitHub repository. Source code for the `CellSpliceNet` model and inference scripts are available under an open-source license at https://github.com/KrishnaswamyLab/CellSpliceNet.

## Supplementary Materials

### Embedding of the expression modality

Figure S1 illustrates the workflow employed to embed the splicing regulatory context from single-cell RNA-seq data using geometric scattering. Initially, we obtain the raw expression count matrix, which is imputed using MAGIC to fill missing values. Next, mutual information is computed pairwise among 243 curated splicing factors, producing discretized co-expression matrices stratified by neuron type. These matrices define co-expression graphs, with nodes representing splicing factors and edges indicating significant mutual dependencies. Finally, we apply the geometric scattering transform on these graphs to generate multiscale, permutation-invariant representations. Here, following the construction of the RNA-secondary-structure and splice-factor co-expression graphs, we apply the *geometric scattering transform* [18] to extract multiscale, permutation-invariant features. For a graph described by adjacency matrix *A* and degree matrix *D*, the lazy diffusion operator is given by

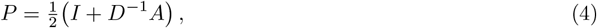

and graph wavelets at scale j are defined as

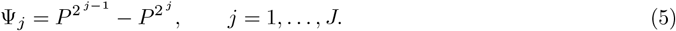

Given a node signal *x* ∈ ℝ^|*V* |^, scattering moments

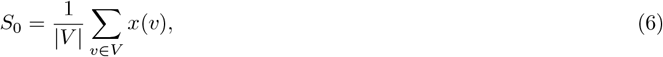

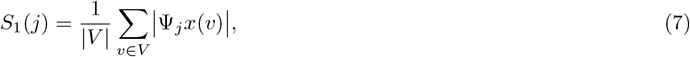

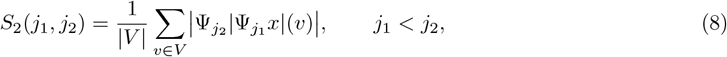

are concatenated for all *j ≤ J* (we use *J* = 4, which gives 11 scales *S*) to yield a fixed-length feature tensor **C** ∈ ℝ^|*V* |×*S*×*d*^.

### Attention in whole RNA sequences

Here we deepen the evidence for the conclusions drawn from Figure 3 of the main manuscript. Across a larger panel of events, they show that (i) `CellSpliceNet` consistently concentrates its attention on intron–exon boundaries, (ii) the precise distribution of those weights is flexibly re-tuned in a neuron-specific, event-dependent manner, and (iii) these attendant differences are captured by low-dimensional PHATE embeddings that reliably cluster neuron classes with similar regulatory signatures. Together, Supplementary Figures S2, S3, S4 demonstrate that the balance between universal splice-site recognition and context-aware modulation is a pervasive property of `CellSpliceNet` rather than an artefact of the examples chosen for the main text.

### Attention in the structural modality

Supplementary Figures S5 and S6 extend the structural-modality analysis of Fig. 4 in the main text. The additional heat-map panels chart nucleotide-level attention across twenty further neurons, reaffirming that `CellSpliceNet` funnels salience toward the exonic midpoint and tapers sharply into the flanking introns (Fig. S5). Overlaying the attention scores on the secondary structures for twenty extra cassette-exon events (Fig. S6) reveals a consistent topological bias: single-stranded loop nucleotides inside the exon draw the highest weights, whereas paired stem regions remain comparatively muted. Together, these expanded examples demonstrate the robustness of `CellSpliceNet`‘s preference for accessible, centrally located motifs when inferring PSI.

**Fig. S1:**
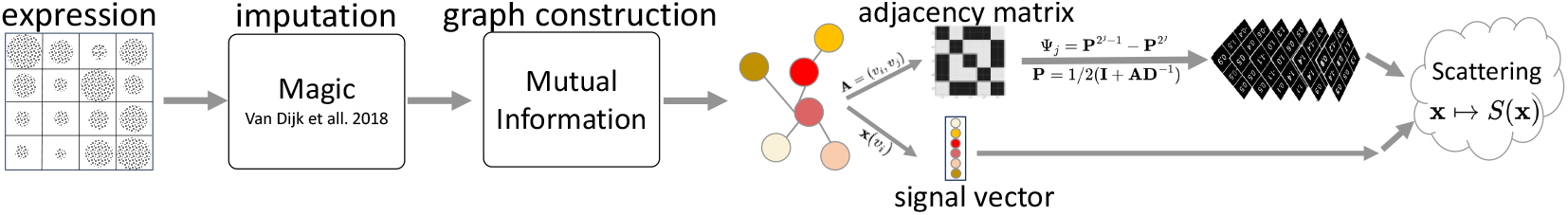
Embedding splicing regulatory context using MAGIC imputation, mutual information-based graph construction, and geometric scattering transform.

### Attention in the ROI modality

We also extend the ROI-level interpretability analysis presented in Fig. 5 of the main manuscript. Fig. S7 aggregates nucleotide-wise attention maps for 20 additional cassette-exon events, confirming that `CellSpliceNet` consistently concentrates weight over the exon core and upstream splice junction while rapidly decaying into the flanking introns. Fig. S8 lifts this view to the motif scale, displaying heat maps in which each subplot corresponds to a neuron type and each row to one of the top ROI motifs.

**Fig. S2:**
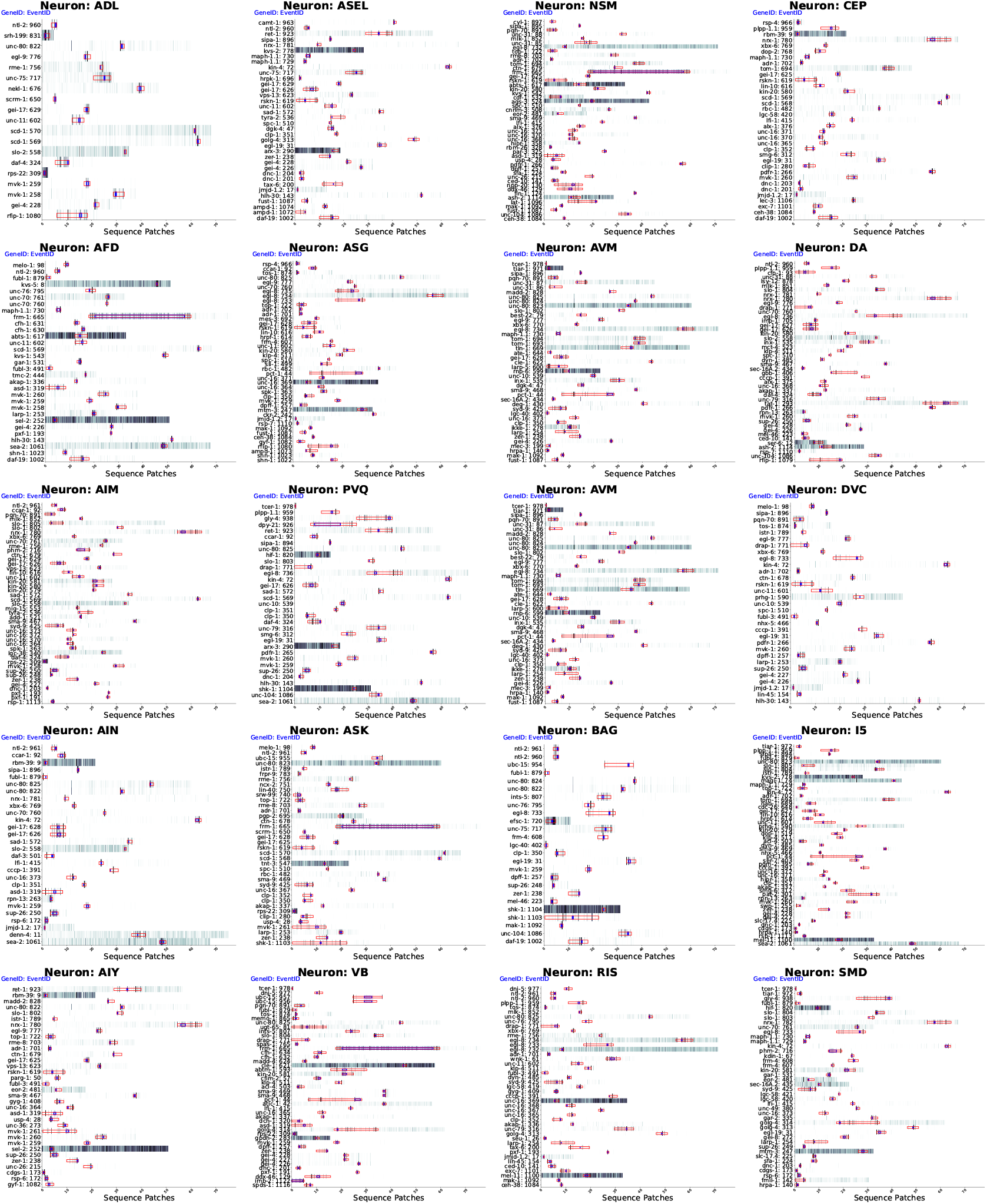
Expanded RNA patch-level attention heat-maps. Heat maps for twenty additional neurons. Color encodes the mean attention weight assigned by `CellSpliceNet`. The blue boxes show the exon regions, and the red boxes show the introns.

**Fig. S3:**
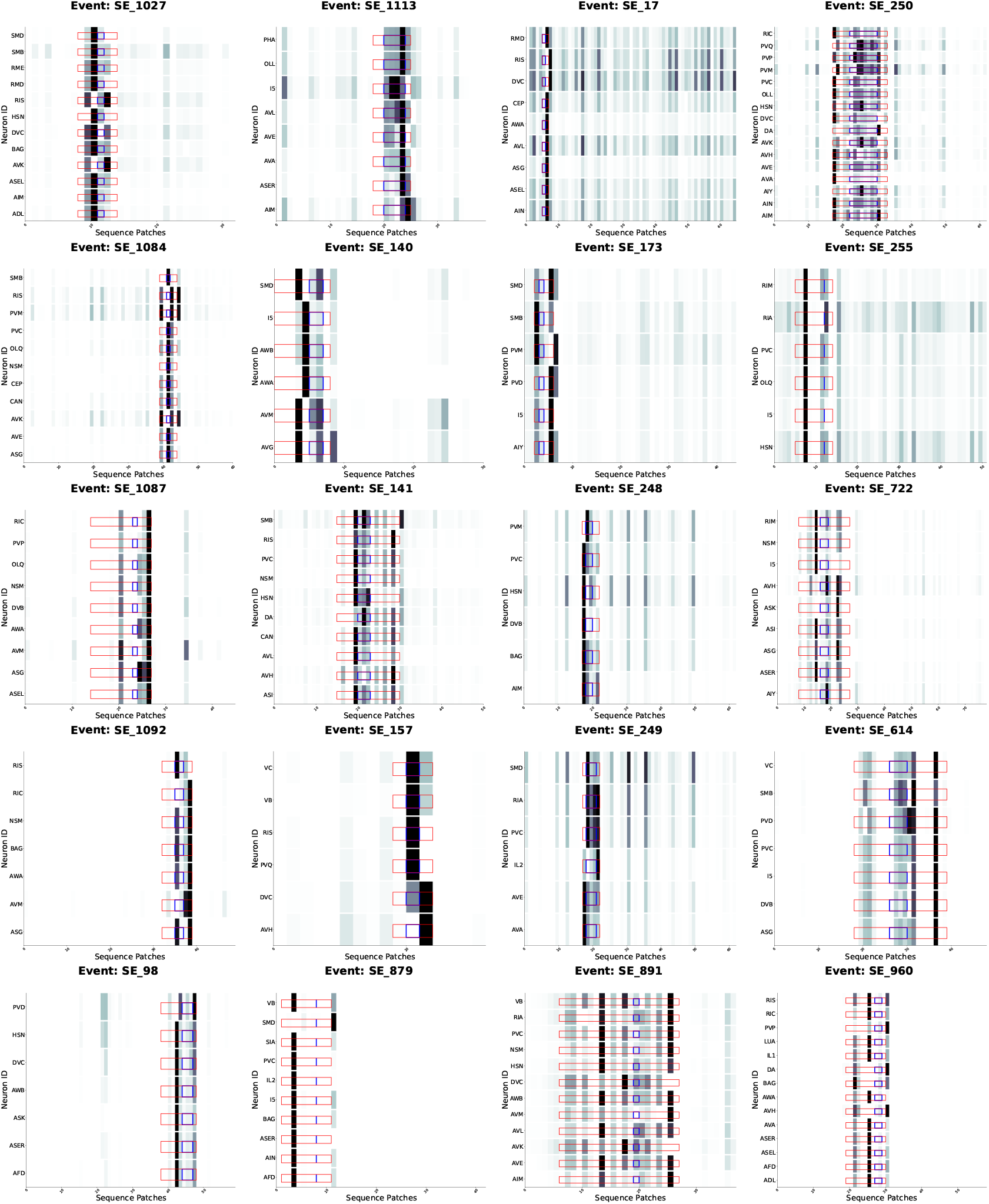
Neuron-specific redistribution of attention across additional events. Stacked bar plots compare per-patch attention profiles for 20 unique events across different neuron types (in each subplot) in each subplot’s y axis.

**Fig. S4:**
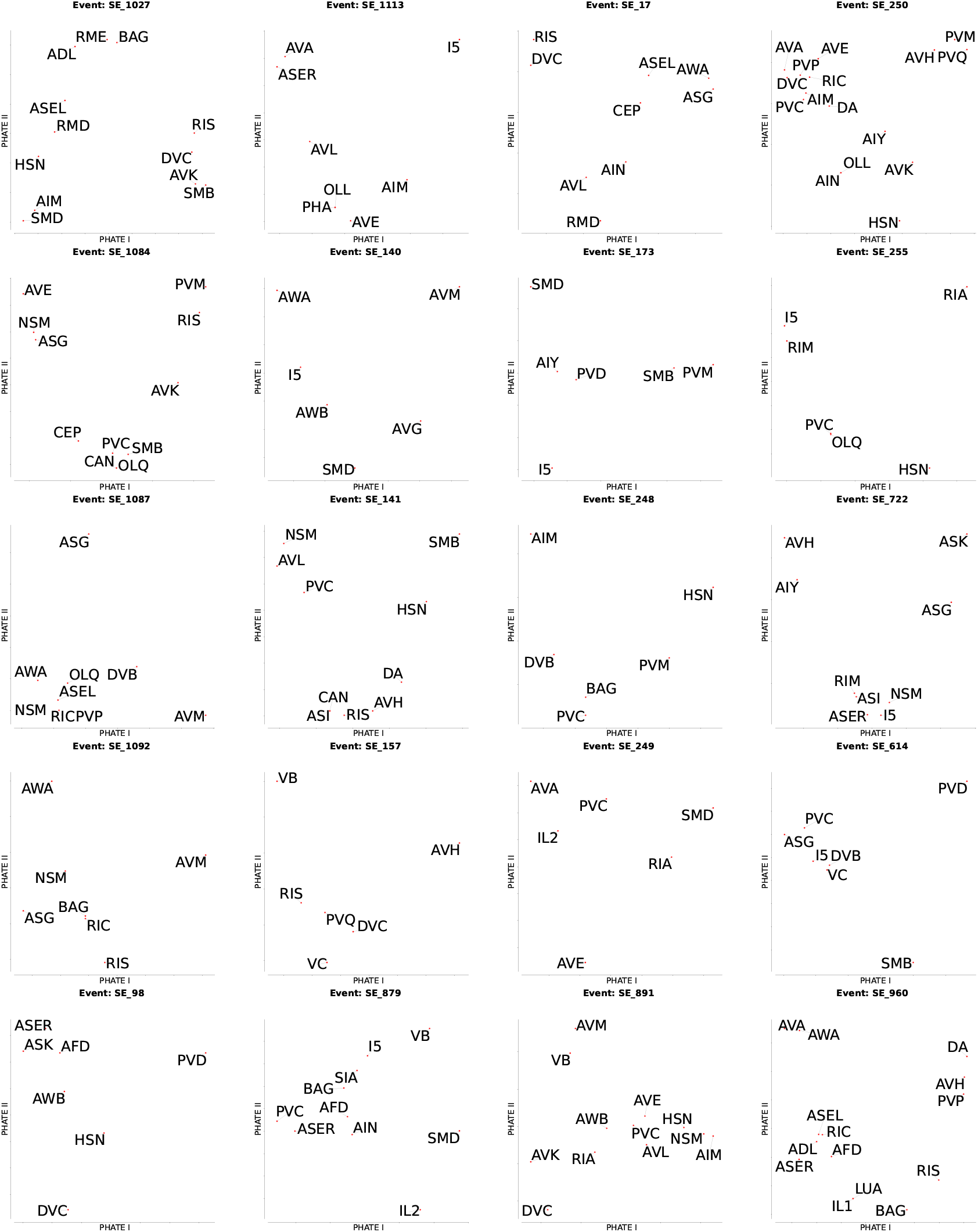
PHATE embedding different events across the neuron types. Each point in a subplot represents a neuron class; axes are the first two PHATE dimensions computed from its attention vector for the corresponding splicing event. Neuron classes with similar attention distributions cluster tightly.

**Fig. S5:**
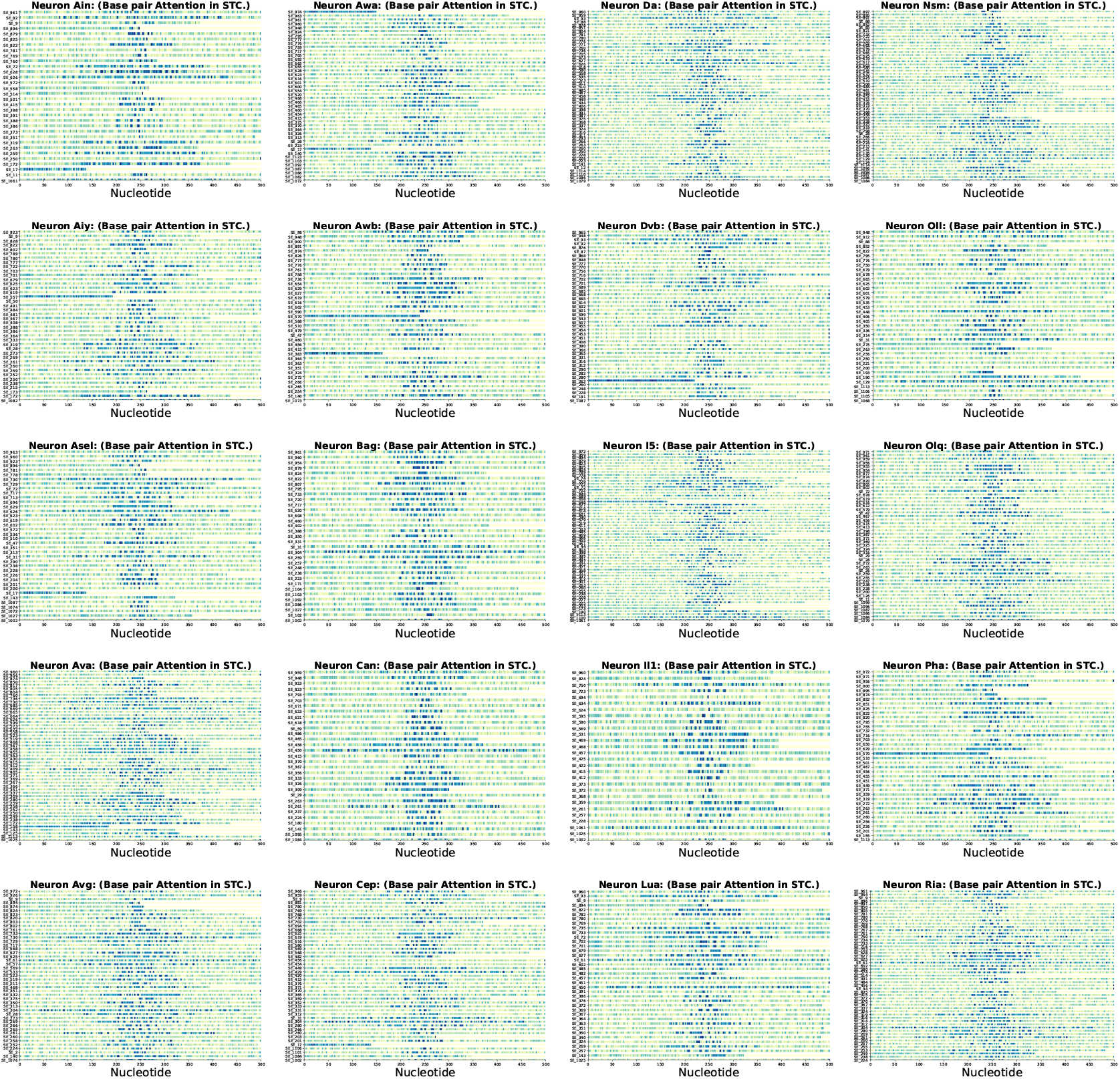
Structure-modality attention concentrates on exonic loop nucleotides. Heat maps depict nucleotide-level attention weights assigned by the structural modality of `CellSpliceNet`. Warmer colors indicate higher salience. Across all cases, attention peaks sharply within the exon center and decays monotonically toward splice junctions and intronic regions.

**Fig. S6:**
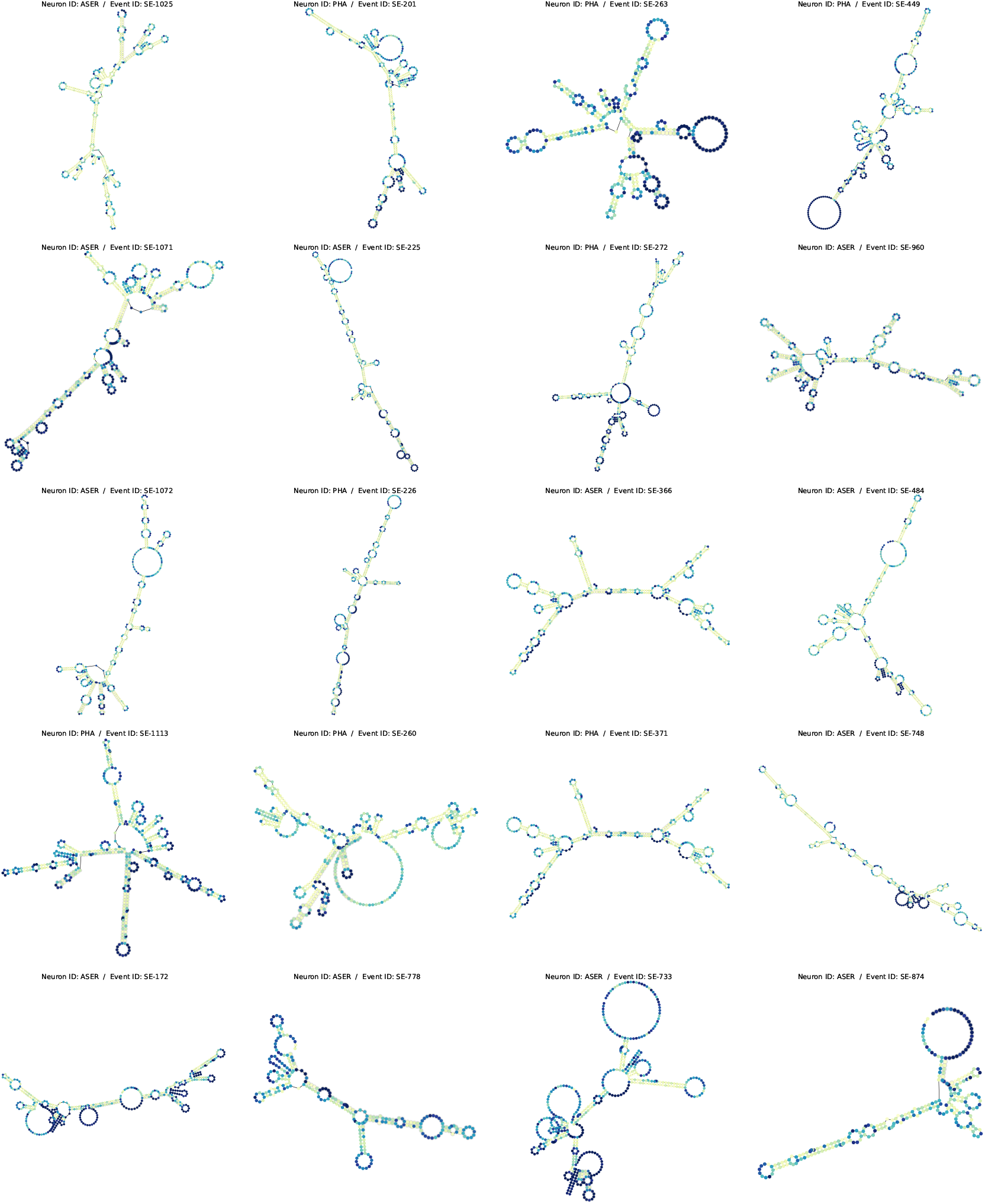
Structure-modality attention concentrates on secondary structures. Attention scores are overlaid on the secondary structures for twenty additional cassette-exon events; dark blue denotes high attention, whereas light yellow indicates low attention as inferred by `CellSpliceNet`.

**Fig. S7:**
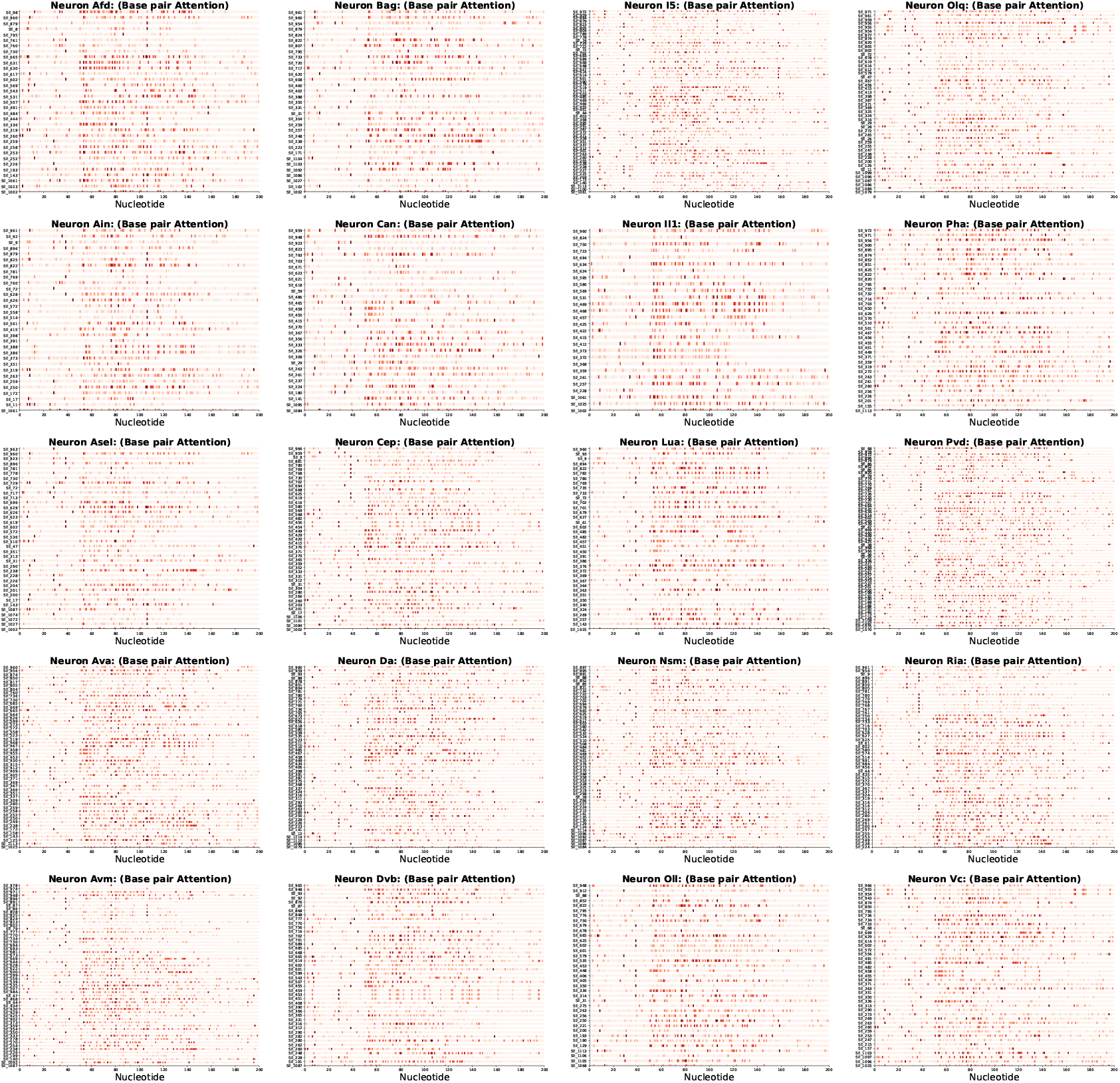
Heightened attention at ROI. Nucleotide-resolution attention maps for representative events of 20 different neuron classes reveal a pronounced focus on the upstream splice boundary.

**Fig. S8:**
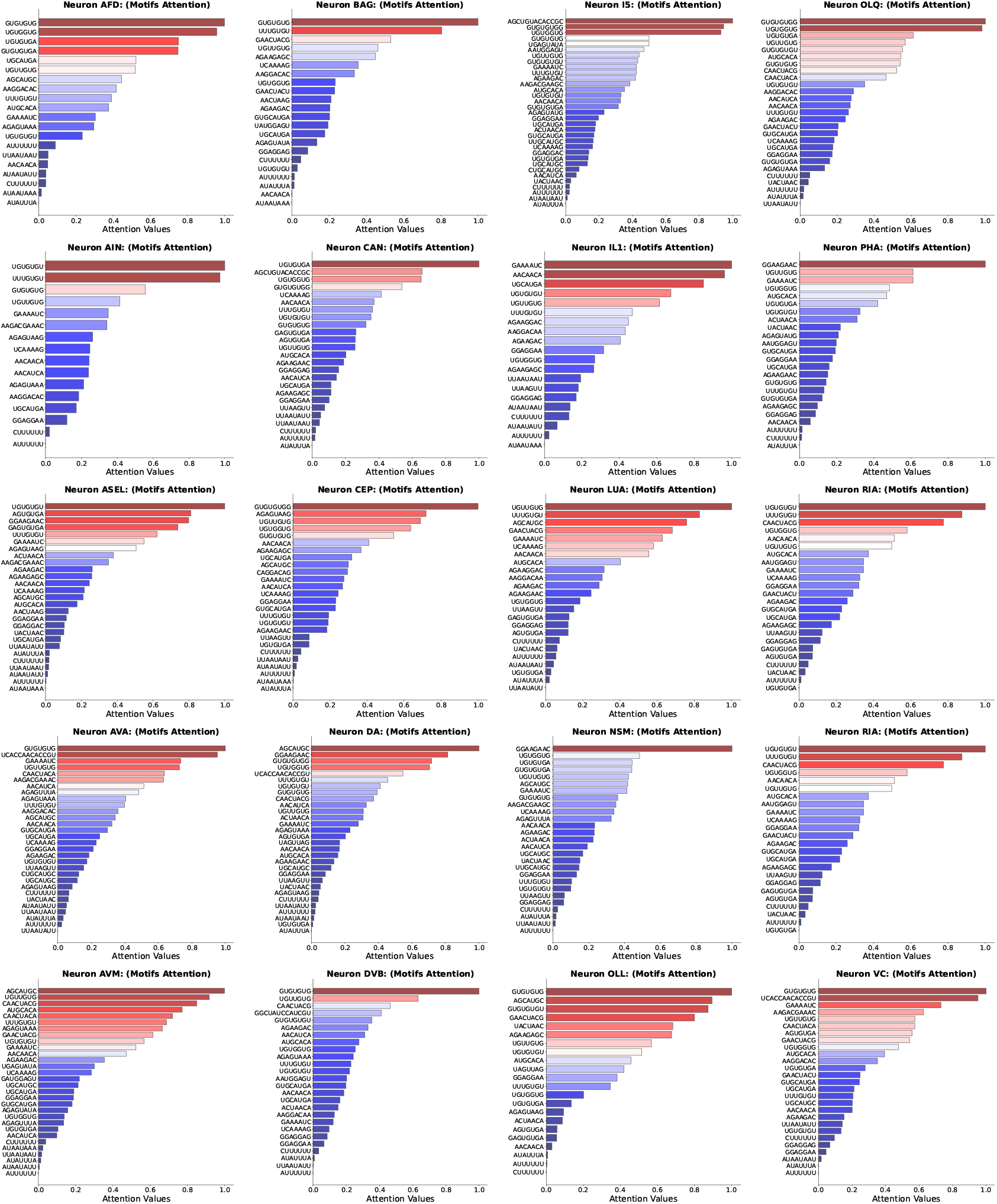
Top motif attention across neuron classes. Motif-level attention is stratified by neuronal type, revealing distinct, class-specific splicing codes.

